# Logic Synthesis of Recombinase-Based Genetic Circuits

**DOI:** 10.1101/088930

**Authors:** Tai-Yin Chiu, Jie-Hong R. Jiang

**Affiliations:** Graduate Institute of Electronics Engineering, National Taiwan University, Taipei 10617, Taiwan.; Department of Electrical Engineering and the Graduate Institute of Electronics Engineering, National Taiwan University, Taipei 10617, Taiwan.

## Abstract

A synthetic approach to biology is a promising technique for various applications. Recent advancements have demonstrated the feasibility of constructing synthetic two-input logic gates in *Escherichia coli* cells with long-term memory based on DNA inversion induced by recombinases. On the other hand, recent evidences indicate that DNA inversion mediated by genome editing tools is possible; powerful genome editing technologies, such as CRISPR-Cas9 systems, have great potential to be exploited to implement large-scale recombinase-based circuits. What remains unclear is how to construct arbitrary Boolean functions based on these emerging technologies. In this paper, we lay the theoretical foundation formalizing the connection between recombinase-based genetic circuits and Boolean functions. It enables systematic construction of any given Boolean function using recombinase-based logic gates. We further develop a methodology leveraging existing electronic design automation (EDA) tools to automate the synthesis of complex recombinase-based genetic circuits with respect to area and delay optimization. Experimental results demonstrate the feasibility of our proposed method.

## I. INTRODUCTION

THE development of synthetic biology shows the feasibility to implement computing devices with DNA genetic circuits in living cells. Synthetic cellular designs often intended to implement certain functions that make cells respond to specific environmental stimuli or even change their growth and cellular development. For instance, synthetic toggle switches [1] and genetic oscillators [2]–[5] can be used to control cell metabolism, synthetic counters [6] can be potentially applied to the regulation of telomere length and cell aggregation, and genetic logic gates [7]–[10] can achieve digital computation in response to stimulus input signals. In addition to these transcription-based DNA circuits, new emerging translational mRNA circuits [11] are likely to have impact on mammalian regenerative medicine and gene therapy. Through the genetic engineering, synthetic cellular circuits are potentially useful to perform therapeutic and diagnostic functions.

For some situations where noxious chemical stimuli exist for many cell generations, the computational results from the synthetic circuits in parent cells are required to be propagated to their daughter cells so that the daughter cells can save time to respond to the environmental stimuli. To achieve this transgenerational memory, one possible method is to store the computational results in separate synthetic memory devices which can be duplicated in cell divisions. In recent work [12], a more efficient scheme for constructing synthetic cellular circuits with integrated logic and memory was proposed, where the computational result was automatically stored in the computing circuit configuration and the changes of configuration can be propagated to its descendant cells. The so-implemented circuits were built based on recombinases and tested in *Escherichia coli* cells and they showed a long-term memory for at least 90 cell generations. More recently, recombinase-based logic circuits has been applied in clinical uses. E.g., in [13] the authors demonstrate that biosensor made of recombinase-based logic gates can be used to detect pathological glycosuria in urine from diabetic patients. The ability to build complex recombinase-based logic circuits is an important step to enable widespread biomedical applications.

Specifically the synthetic cellular circuits proposed in [12] used serine recombinases Bxb1 and phiC31 to implement various two-input logic gates. A serine recombinase targeting a pair of non-identical recognition sites known as *attB* (**att**achment site **b**acteria) and *attP* (**att**achment site **p**hage) is able to induce irreversible DNA inversion. As illustrated in Fig. 1(A), since the inversion makes the recognition sites become hybrid sites called *attR* and *attL* which cannot be targeted by the recombinase, no further inversion is allowed afterwards.

**Fig. 1.**
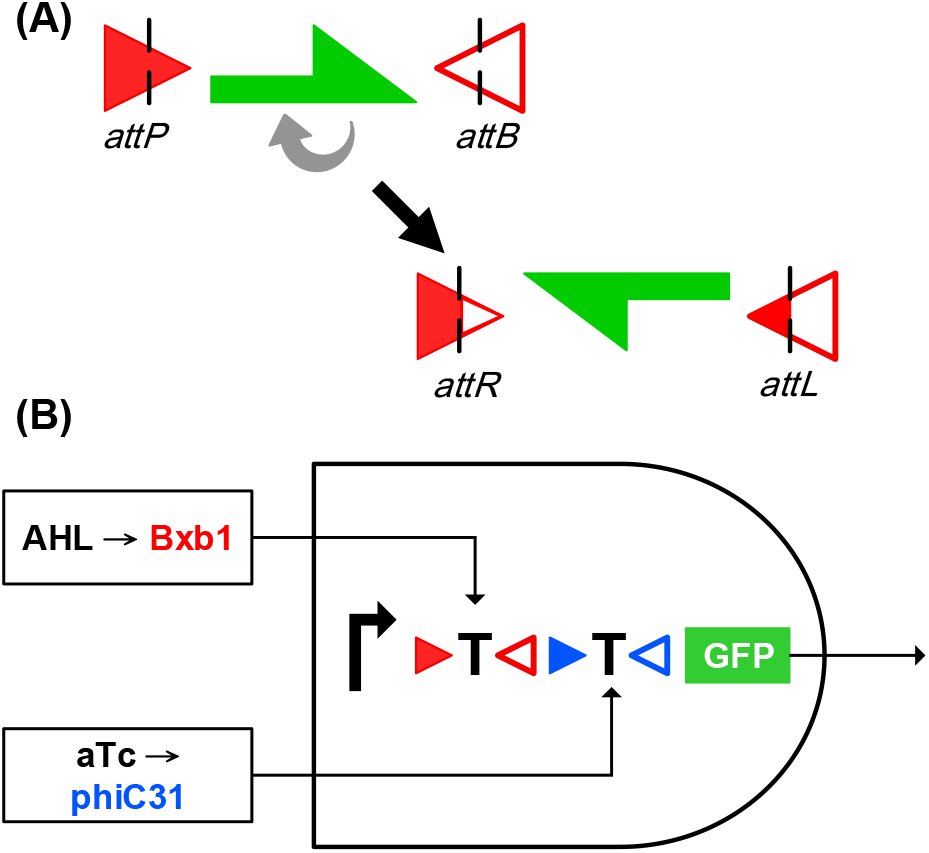
(A) Schematic illustration of the irreversible inversion of DNA sequences using serine recombinases. (B) Implementation of an AND gate using recombinases. The right-turn arrow represents a promoter; the red and blue triangles are the targeting sites of recombinases Bxb1 and phiC31, respectively; the letter T’s flanked by the targeting sites are transcription terminators; the green box represents the gene encoding the green fluorescent protein.

We illustrate how recombinases take part in the implementation of two-input logic gates with the two-input AND gate example shown in Fig. 1(B). (As a convention, in this paper we read a DNA sequence from left to right assuming the 5’-to-3’ direction of the coding strand.) Let molecules AHL and aTc be the stimulus inputs to a cell and act as inducers activating the expressions of recombinases Bxb1 and phiC31, respectively. These recombinases when activated will irreversibly invert (flip) the DNA sequences flanked by their recognition sites (denoted by the colored triangle pairs). The DNA sequences being flanked can be a promoter, a transcription terminator, or a reporter, e.g., a green fluorescent protein (GFP). Inverting these DNA sequences will alter the output gene expression. In Fig. 1(B), two terminators were flanked by the recognition sites of recombinases Bxb1 and phiC31, and the output green fluorescent reporter is highly expressed only when both inducers AHL and aTc are in high concentration to activate BxB1 and phiC31 which together further flip and disable both terminators (denoted by letter “T”). Therefore, the circuit of Fig. 1(B) effectively implements a two-input AND gate. Note that such DNA sequence changes will survive through cell divisions and can be inherited to descendant cells in different generations. Hence the so-implemented logic function can achieve a long-term transgeneration memory.

Note that the feasibility of constructing large recombinase-based circuits is limited to available recombinases. Nevertheless, with the advances of biotechnology, DNA inversion techniques mediated by genome editing approaches, such as ZFNs [14], [15], TALENs [15]–[17], and CRISPR-Cas9 nucleases [17]–[20] have already been reported. It is envisaged that these genome editing tools could be alternatives scalable to realize large recombinase-based circuits [21]. Motivated by the viability and applicability of recombinase-based circuits, in this paper we formalize the construction of a general multi-input logic gate with its DNA sequence composed of series of promoters and transcription terminators targeted by multiple recombinases. We further characterize the set of Boolean functions realizable under such logic gates. In addition, we show a design flow for arbitrary Boolean function construction with cascaded recombinase-based logic gates. This automated design methodology is demonstrated by leveraging synthesis tool ABC [22], an electronic design automation (EDA) tool developed at UC Berkeley, to synthesize cascaded multi-level recombinase-based circuits.

The rest of the paper is organized as follows. In Section II, some examples of multi-input recombinase-based logic gates are shown to motivate this work. In Section III, the syntax and semantics of recombinase-based logic gates are formalized. In Section IV, we propose a method to synthesize logic circuits composed of recombinase-based gates using conventional logic synthesis tools. In Section V, experimental results are evaluated. Finally, conclusions and future work are remarked in Section VI.

## II. PRELIMINARIES

To formalize the general multi-input gate construction, we use the three-input logic gates in Fig. 2 as an example to illustrate. Fig. 2(A) shows a realization of a 3-input AND gate using three recombinases *R*_*1*_, *R*_*2*_, and *R*_*3*_, where molecule *I*_*i*_is a stimulus input that activates the expression ofrecombinase *R*_*i*_, for *i* = 1, 2, 3. Then *R*_*i*_*’*s induce the inversions of their corresponding DNA sequence fragments. In order to express GFP in this gate, first we require *R*_*1*_ to invert the inverted promoter so that the RNA polymerase can bind to it and begin the transcription of the downstream DNA sequence in which the GFP gene resides. Second, *R*_*2*_ is needed to flip the terminator to avoid the termination of transcription before reaching the GFP gene. Third, *R*_*3*_ is demanded to upright the GFP gene for the RNA polymerase to initiate GFP production. Collectively, to have GFP highly expressed all *R*_*i*_’s must exist, and thus this circuit implements a 3-input AND gate. Note that this 3-input AND gate, where the promoter and the reporter gene GFP can be flipped by recombinases, is designed in a different fashion from the 2-input AND gate in Fig. 1(B), where only transcription terminators are inverted by recombinases. The additional choice of flipping the DNA fragments of promoter and GFP gives more flexibility for logic gate construction.

**Fig. 2.**
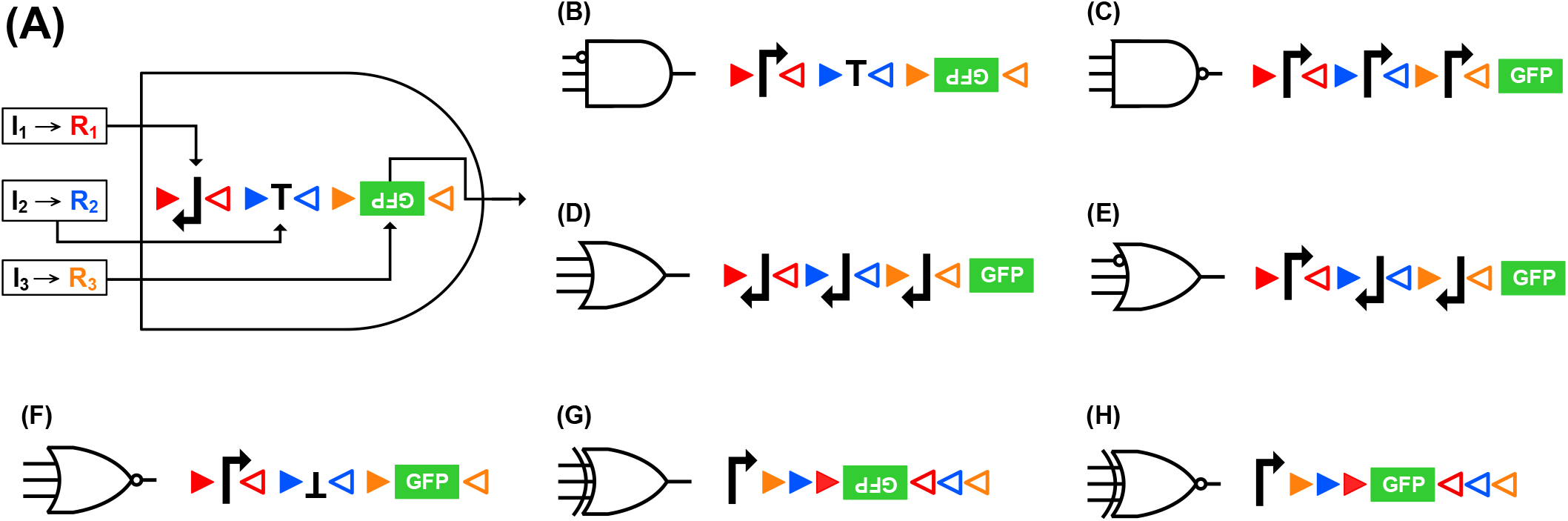
Implementation of basic 3-input logic gates using recombinases. The inputs of each gate from top to down are recombinases *R*_*1*_, *R*_*2*_, and *R*_*3*_, respectively; inducer *I*_*i*_, monitored by the cell activates the expression of *R*_*i*_; the red, blue, and orange triangles denote the targeting sites of *R*_*i*_*, i* = 1, 2, 3, respectively.

In Fig. 2(B)-(H) we present seven other basic 3-input gates implemented with recombinases. Special implementations with nested targeting sites are applied on the XOR gate in (G) and the XNOR gate in (H). In the XOR gate in (G), the existence of one or three recombinases results in one or three times of GFP gene flipping and thus making the upside-down gene become upright, while the existence of two recombinases makes the GFP gene flip twice and remain upside down. Similar situations happen in the XNOR gate in (H).

Since the implementations of multi-input gates are possible, we are not constrained to using only 3-input gates and basic gate types, such as AND, OR, NAND, NOR, XOR, and XNOR gates. Rather, we can construct complex logic gates with more inputs. Fig. 3(A) shows an example of a 4-input logic circuit

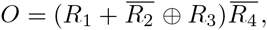

which can be directly realized by a single 4-input complex logic gate as shown in Fig. 3(B), instead of cascading multiple two-input gates.

**Fig. 3.**
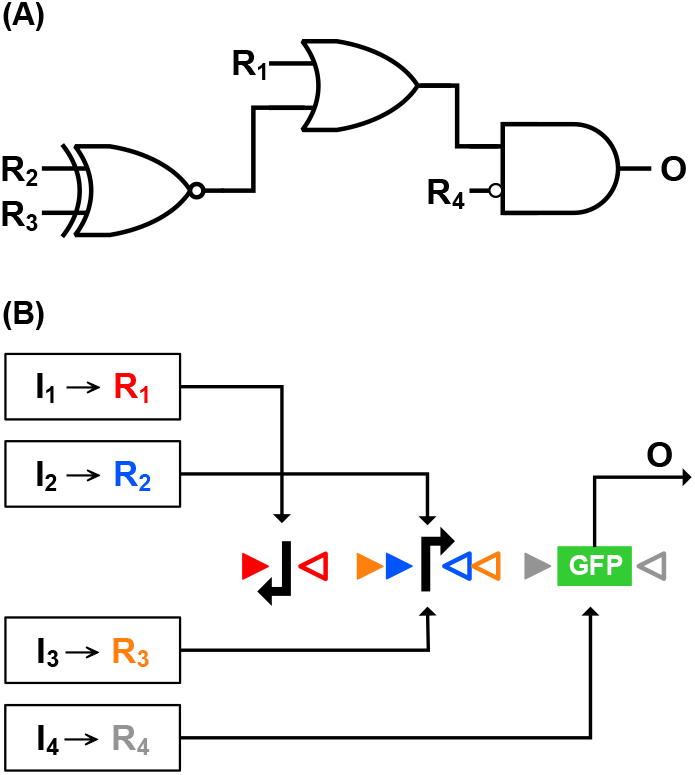
(A) Schematic illustration of a 4-bit non-basic logic function 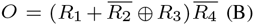 (B) Corresponding implementation using recombinases.

## III. FORMALISM OF RECOMBINASE-BASED LOGIC GATES

### A. Syntax of Well-formed Sequences

We define the following syntax to formalize the DNA sequences of logic gates constructed with recombinases. Here the basic elements composing a legal DNA sequence of a recombinase-based logic gate are “atomic terms,” including (inverted/non-inverted) transcription factors, (inverted/non-inverted) promoters, (inverted/non-inverted) genes, and targeting sites of recombinases. The syntax of DNA sequence forming a legal recombinase-based logic gate can be defined as follows.

*Definition 1:* An *atomic term* in a DNA sequence is a transcription terminator *T*, a promoter *P*, a gene *G*, an inverted transcription terminator 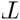, an inverted promoter 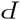, or an inverted gene 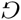. The syntax of an atomic term can be expressed in Backus-Naur Form as

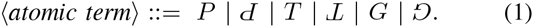

Let the targeting sites *attP* and *attB* of recombinase *r* in a DNA sequence be denoted as 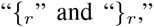 respectively. In the sequel, the subscripts of 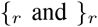 may be omitted for brevity when they are clear from the context or immaterial to the discussion. Note that targeting sites 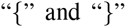 of a recombinase must appear in a pair.

*Definition 2:* The syntax of a *well-formed sequence* (wfs) is recursively defined as follows.

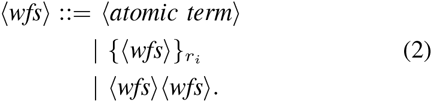

In this paper we concentrate on the special case of one-gene *wfs* (1g-wfs), where only one gene *G*, which is neither inverted nor sandwiched by targeting sites, appears in the wfs at the end of the sequence serving as the output. For example, 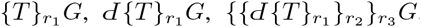, and 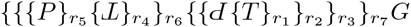 are 1g-wfs’s. Notice that under the 1g-wfs setting, the logic gate has a single output and the gene can only be transcribed in one direction from left to right.

A pair of targeting sites of a recombinase is called *basic* if it only flanks an atomic term. Otherwise, it is called *non-basic*. We call a 1g-wfs *basic* if it contains only basic pairs of targeting sites, and *non-basic* if it contains some non-basic pair of targeting sites. For example, 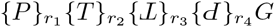 is a basic 1g-wfs. In contrast, 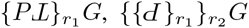, and 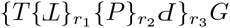 are non-basic 1g-wfs’s.

Furthermore, a non-basic pair of targeting sites can be nested. That is, a non-basic pair of targeting sites can be flanked by another pair of targeting sites. For instance, 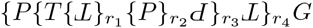 has nested two pairs of targeting sites targeted by the recombinases *r*_*3*_ and *r*_*4*_.

We discuss the logic functions induced by basic and non-basic 1g-wfs’s in the following.

### B. Semantics of Well-Formed Sequences

#### 1) Basic well-formed sequences

We first study some reduction rules of basic 1g-wfs’s. Let *σ* be the DNA sequence of a basic 1g-wfs excluding the output gene, that is, *σ* is a basic wfs without any gene. We denote a wfs without any gene as 0g-wfs. Because *σ* is made of components P, 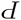, T, 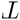, 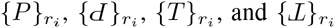 for any component *C* in *σ*, the sequence *σ* can be decomposed into

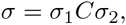

where *σ*_1_ and *σ*_2_ are two 0g-wfs’s, if non-empty. We show that the logic gate induced by the 1g-wfs *σG* can be further reduced to an equivalent form according to the type of the component *C*.

When *C* is a transcription terminator *T*, then *σ* equals 
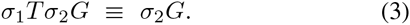

This equivalence holds because any transcription that starts from *σ*_1_ to gene *G* is always blocked by the transcription terminator *T* in the middle, making *σ*_1_*T* a don’t-care and thus removable.

When *C* is an inverted terminator 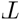, then *σ* equals

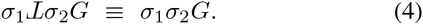

This equivalence holds because the inverted terminator 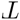 never blocks the transcription and is thus removable.

When *C* is a promoter *P*, then *σ* equals

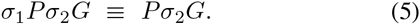

This equivalence holds because no matter whether there is a transcription that starts from *σ*_1_ to *G* or not, a transcription can always start from the promoter *P*. Therefore, *σ*_1_ is a don’t-care and thus removable.

When *C* is an inverted promoter 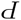, then *σ* equals

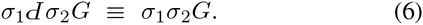

This equivalence holds because the transcription that begins at 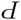 proceeds across *σ*_1_ in the direction from right to left, it does not pass through *G*. As a result, the expression of *G* can not be initiated by 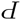 and thus 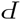 can be removed from the sequence.

When *C* is 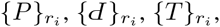, or 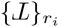, since an atomic term *A* is equivalent to {*A*}_*r*_ for recombinase *r* being in low concentration (denoted *R* = 0 by treating *r* as a Boolean variable *R* of value 0) or 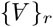 for recombinase *r* being in high concentration (denoted *R* = 1 by treating *r* as a Boolean variable *R* of value 1), the reduction rules for *C* can be easily extended from the previous rules as summarized below.

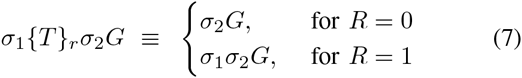

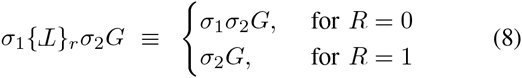

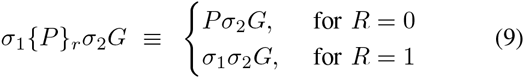

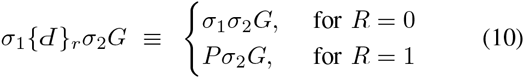

With the above analysis, we can derive the corresponding Boolean function of a given 1g-wfs. Consider the 1g-wfs *σG* with the sequence *σ* targeted by recombinases *r*_*i*_*, i* = 1 ⋯ *n*. Activating the expression of gene *G* requires the recombinases *r*_*i*_*’*s have adequate (high or low) concentrations so that the 1g-wfs *σG* effectively reduces to *PG*. The Boolean function induced by *σG* is determined through a series of decisions made by *r*_*i*_’s. In essence, it corresponds to a decision list [23]. To illustrate, consider the example 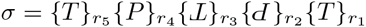. The decision list induced by the 1g-wfs *σG* is shown in Fig. 4. Note that given a sequence without non-basic targeting sites, the decisions always start from the rightmost to the leftmost components because a component closer to the gene may overwrite the effects imposed by the components on its left and thus it is of higher priority. Therefore, the Boolean function of *σG* is determined starting from *R*_*1*_ to *R*_*5*_. In order to reduce *σ* to P to express gene *G*, first we must require *R*_*1*_ to be 1. Otherwise if *R*_*1*_ = 0, *σ* becomes equivalent to a null sequence no matter what other *R*_*i*_’s are. Next, if we let *R*_2_ be 1, we can have an equivalent sequence equal to *P* as wished. Otherwise we can let *R*_2_ be 0 and look for other possibilities for the reduction to *P*. If *R*_2_ = 0, we can easily tell that the only possibility occurs when *R*_3_ and *R*_4_ are both 0 and that the logic of *R*_5_never affects the reduction. Collectively, the logic function of the gate σG is derived as 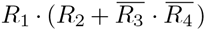, where symbol “+” denotes Boolean disjunction, symbol “•” denotes Boolean conjunction, and symbol “-” or “!” denotes Boolean negation. In the sequel, we sometimes omit the conjunction symbol “•” in a Boolean expression.

**Fig. 4.**
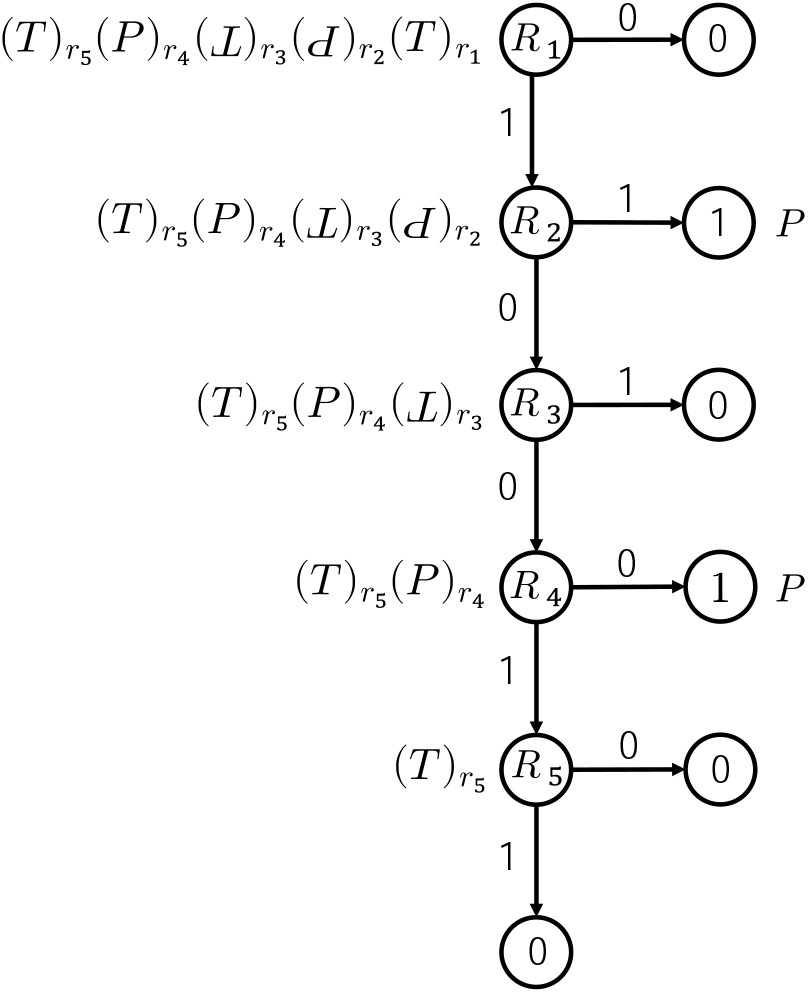
Decision list corresponding to 1g-wfs 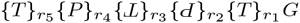. Node labelled *R*_*i*_ is the decision for the logic value of Nodes labelled 0 (resp. 1) stand for gene *G* cannot (resp. can) be expressed. The sequences beside nodes are the equivalent sequences after the corresponding (partial) decisions.

In general, we can systematically convert any basic 1g-wfs to its corresponding logic function. To achieve this conversion, the operator Ω over a 1g-wfs is defined in Table I. For an empty sequence ⊥, we define Ω[⊥] = 0. E.g., for the 1g-wfs 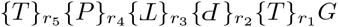, its Boolean function is derived by

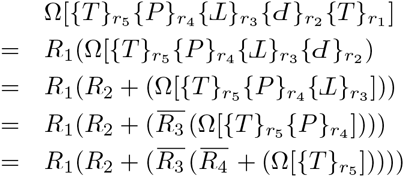

**TABLE I.** Operators for Parsing Basic 1G-WFS *σCG*, with (Non-Empty) 0G-WFS *σ*, Component *C*, and Gene *G*, to Logic Function.

| component $C$ | operator $\Omega[\sigma C]$ |
| --- | --- |
| $T$ | $0 \cdot (\Omega[\sigma])$ |
| $P$ | $1 + (\Omega[\sigma])$ |
| $\{T\}_r$ | $R \cdot (\Omega[\sigma])$ |
| $\{P\}_r$ | $\bar{R} + (\Omega[\sigma])$ |
| $\mathcal{L}$ | $1 \cdot (\Omega[\sigma])$ |
| $\mathcal{d}$ | $0 + (\Omega[\sigma])$ |
| $\{\mathcal{L}\}_r$ | $\bar{R} \cdot (\Omega[\sigma])$ |
| $\{\mathcal{d}\}_r$ | $R + (\Omega[\sigma])$ |

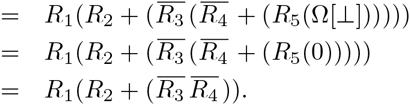

#### 2) Non-basic well-formed sequences

We extend the above derivation of Boolean function to non-basic 1g-wfs’s by having the operator Ω over a 0g-wfs {σ}_*r*_ (which can be basic or non-basic) defined as

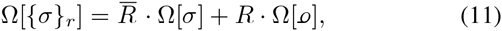

where 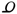 is the inverted sequence of *σ*. To understand Eq. (11), consider a 1g-wfs *σG* with only one pair of non-basic targeting sites. Suppose 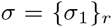, where *σ*_1_ is a basic 0g-wfs. Then σ is equal to *σ*_1_ when *R* =0 and to 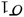, the inverted sequence of *σ*_1_, when *R* =1. For example, the logic function for 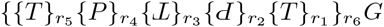 can be obtained by

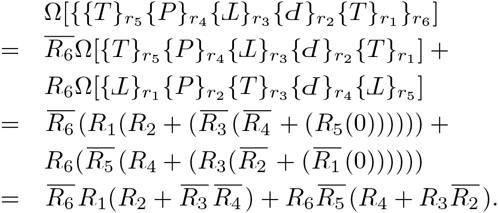

For a 1g-wfs with multiple (possibly nested) non-basic pairs of targeting sites, its logic function can also be directly derived by the Ω operator. For example, the logic function for 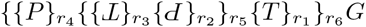 can be obtained by

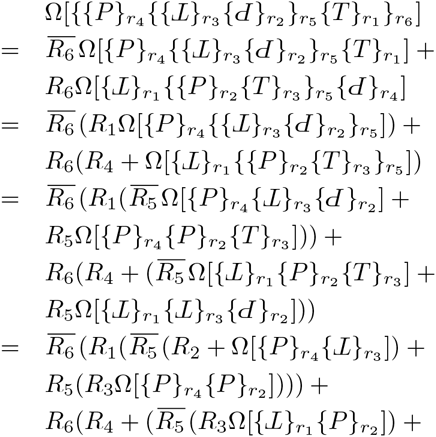

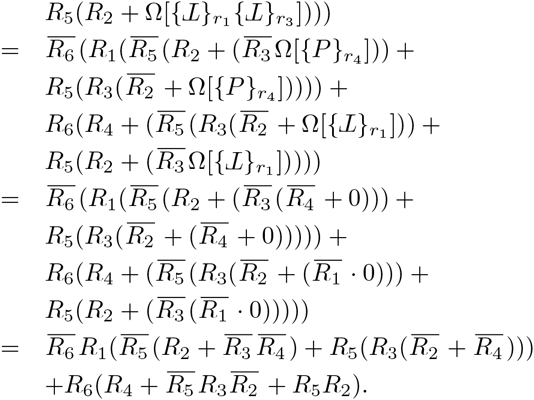

Non-basic pairs of targeting sites can be exploited to efficiently construct special Boolean functions. One of such special functions is the parity function. An *n*-input odd parity function can be realized by the 1g-wfs

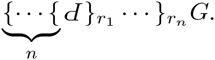

When there is an odd number of *R*_*i*_’s equal to 1, the 1g-wfs reduces to sequence *PG* and gene *G* can be expressed. Otherwise it reduces to sequence G and gene G cannot be expressed. On the other hand, the *n*-input even parity function can be realized by the 1g-wfs

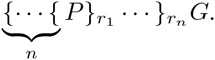

Note that in the formation rule of well-formed sequences in Eq. 2, a pair of targeting sites appears inductively. A DNA sequence, e.g., 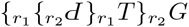 shown in Fig. 5, with interlocking pairs of targeting sites is not included in the definition of Eq. (2). Such a sequence is excluded due to the fact that the inversions caused by interlocking pairs of targeting sites may result in nondeterministic behavior. For example, in Fig. 5 the expression of *GFP* is nondeterministic when both *r*_1_ and *r*_2_ are of high concentrations. If recombinase *r*_1_ inverts the sequence flanked by the red triangles first, the terminator *T* can no longer be inverted by recombinase *r*_2_, and thus *GFP* cannot be expressed. In contrast, if the inversion is made by recombinase *r*_2_ first, then *GFP* can be expressed. Depending on which recombinase acts first, the output of *GFP* is nondeterministic. Although sequences with interlocking pairs of targeting sites can exhibit interesting nondeterministic behaviors with memory, how to construct systems with such sequences is out of the scope of this work.

**Fig. 5.**
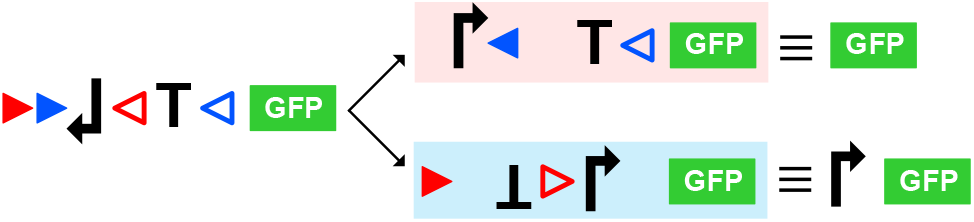
A 1g-wfs with two pairs of targeting sites interlocking with each other. The red and blue pairs denote the targeting sites of recombinase *r*_1_ and *r*_2_, respectively. The effective 1g-wfs’s after the inversions induced by *r*_1_ and *r*_2_ are shown in the red and blue panels, respectively, which are followed by their equivalent simplified sequences.

## IV. CONSTRUCTION OF MULTI-LEVEL RECOMBINASE-BASED LOGIC CIRCUITS

With the recombinase-based logic gates built from 1g-wfs’s, we can cascade them to implement arbitrary complex multi-level circuits. For example, the logic function 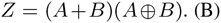 can be implemented with the two-level circuit shown in Fig. 6(A), which is composed of an OR-gate, an XOR-gate, and an AND-gate. One possible DNA implementation of *Z* with cascade can be derived by converting each gate to their 1g-wfs realizations as shown in Fig. 6(B). The 1g-wfs’s that encode the genes *R*_1_, *R*_2_, and *Z* correspond to the OR, XOR and AND gates, respectively. The recombinases *r*_1_ and *r*_2_ as the inputs to the AND gate are the intermediate signals.

**Fig. 6.**
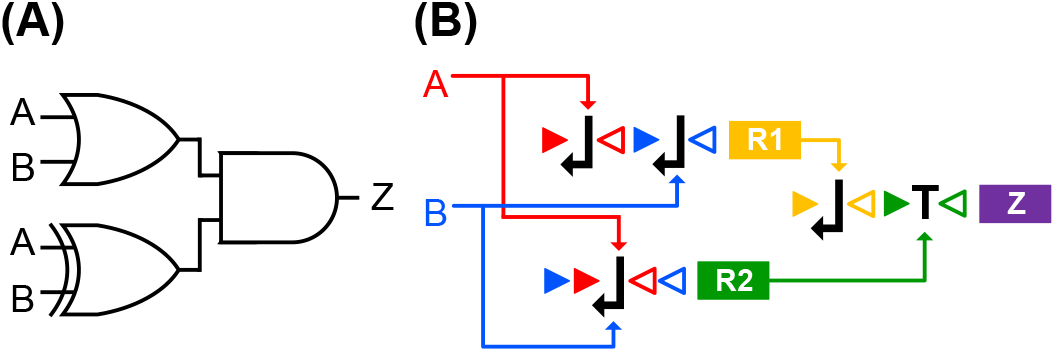
(A) Logic circuit of Boolean function 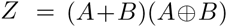. The corresponding DNA implementation of the circuit in (A) with gate cascade. *A* and *B* denote the recombinase inputs of the overall circuit. The genes *R*1 and *R*2 encode the recombinases *r*_1_ and *r*_2_, respectively, which are the inputs to the downstream AND gate. The protein encoded by the gene *Z* is the output of the circuit.

Because the basic 1g-wfs gates can implement decision list functions, they form a *functionally complete* set of primitive logic gates that can be composed to implement any Boolean function. Therefore the 1g-wfs gates can be collected as a library for the synthesis of complex logic circuits. By leveraging conventional logic synthesis tools in electronic design automation (EDA), recombinase-based logic circuits can be synthesized with the flow shown in Fig. 7. Given a Boolean function or circuit netlist as the input, it is first optimized by technology-independent techniques for circuit simplification. The simplified circuit is further optimized by technology-dependent techniques for technology mapping using the primitive gates in the given standard cell library. To achieve recombinase-based logic circuit synthesis, the main task is to provide the library while all other optimization tasks can be done using existing logic synthesis tools.

**Fig. 7.**
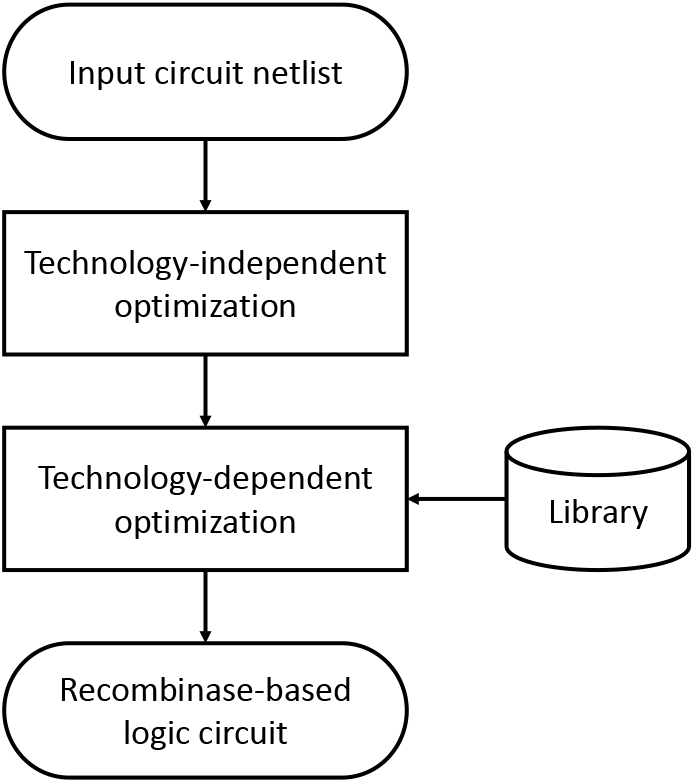
Logic synthesis flow for the implementation of recombinase-based logic circuit

In this work, we adopt ABC [22], an industrial-strength logic synthesis tool developed at UC Berkeley, for circuit synthesis and optimization. Given a circuit netlist, we first apply ABC to perform technology-independent optimization on the netlist, e.g., Boolean minimization to minimize the number of product terms and literals. We then use ABC to perform technology mapping to implement the area or performance optimized netlist using the 1g-wfs gates in the library.

To illustrate the synthesis flow, we consider implementing ISCAS benchmark circuit c17 shown in Fig. 8 with recombinase-based genetic circuit realization. The circuit consists of five inputs *A*, *B*, *C*, *D*, and *E*, and two outputs *Y* and *Z* with functions

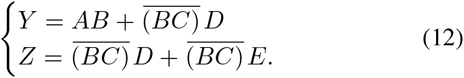

**Fig. 8.**
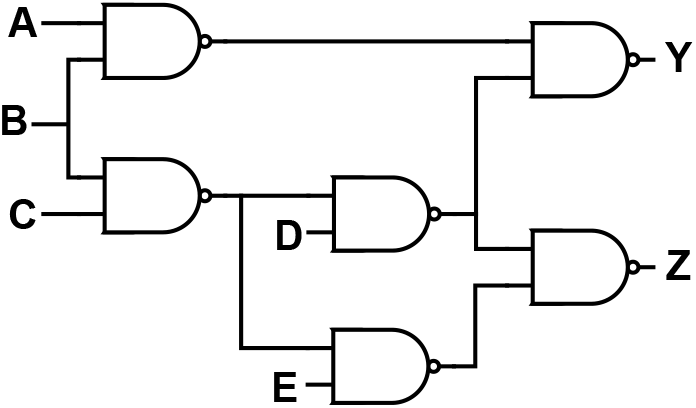
Circuit diagram of ISCAS benchmark c17. c17 circuit consists of six NAND gates with five inputs {*A*, *B*, *C*, *D*, *E*} and two outputs {*Y*, *Z*}.

For area-driven synthesis of benchmark c17, there are 44 DNA gates defined by their 1g-wfs’s with up to three recombinase inputs. They are collected as the library as shown in Fig. 9. According to the experiment in [12], where the promoters and transcription terminators used are roughly of the same length, we treat the area cost of both promoter and transcription terminator as unity. Therefore, the area cost of a DNA gate is defined as the number of atomic terms, excluding the output gene, that appear in the 1g-wfs of the gate. For example, the gate c3_1 corresponding to a 3-input OR gate has three inverted promoters as shown in Fig. 2(D). Hence, the area cost of c3_1 is counted as 3 units. By providing the c17 netlist and the library to ABC, the tool can perform optimization and technology mapping to find an area-optimized circuit composed of DNA gates of the library.

**Fig. 9.**
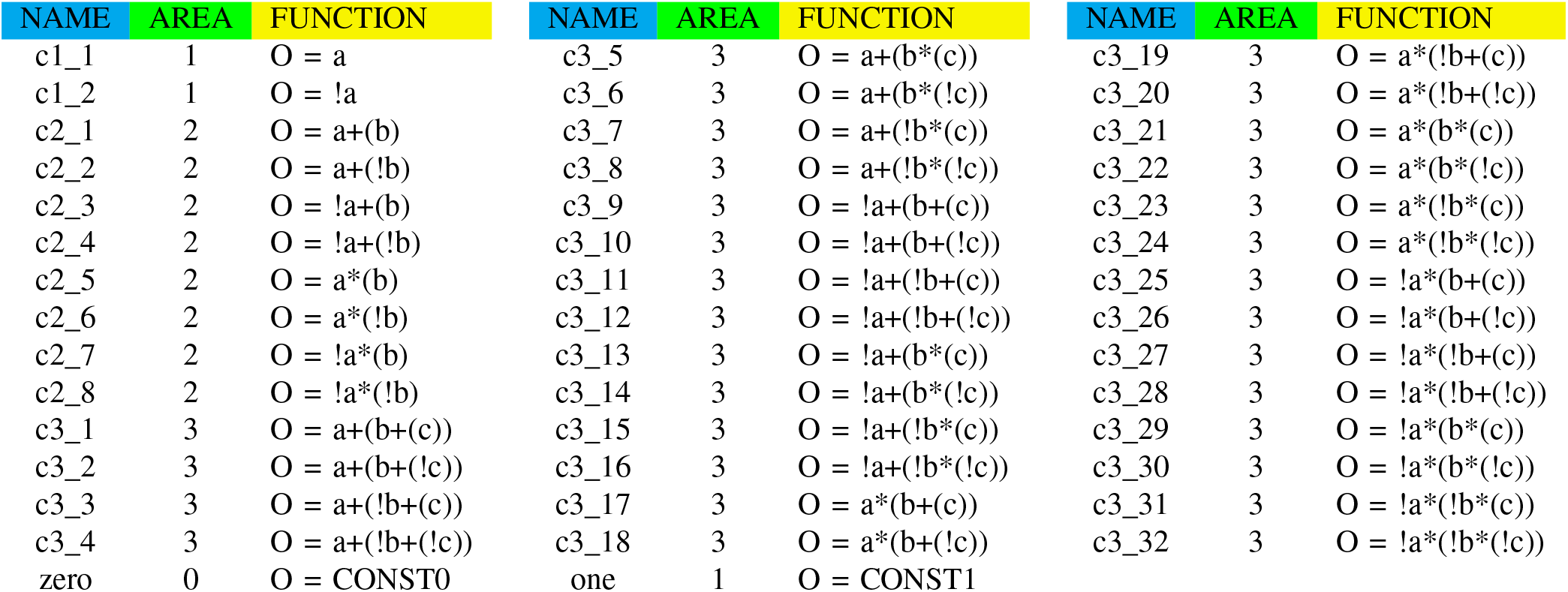
Library of DNA gates with specification of area cost. The library contains 44 different cells and each cell corresponds to a DNA logic gate defined by a 1g-wfs with up to three inputs. The variables *a*, *b*, and c in a function specification represents the recombinase inputs to a gate, and the variable *O* denotes the gate output.

Fig. 10 shows the result described in Verilog language of the synthesized c17 recombinase-based circuit using library gates listed in Fig. 9. The synthesized circuit comprises gates c2_4, c2_5, c3_14, and c3_25, and the total area cost is 10 units. Note that the naive DNA circuit implementation of c17 circuit by converting the digital logic gates in Fig. 8 to the corresponding DNA gates results in a total area cost of 12 units. Compared to the naive implementation, the area cost of the circuit synthesized by ABC technology mapping decreases. The logic functions of *Y* and *Z* in the synthesized circuit can be easily verified to be consistent with Eq. (12), implying the correctness of the synthesis result. The DNA circuit of module c17 in Fig. 10 is plotted in Fig. 12(A).

**Fig. 10.**
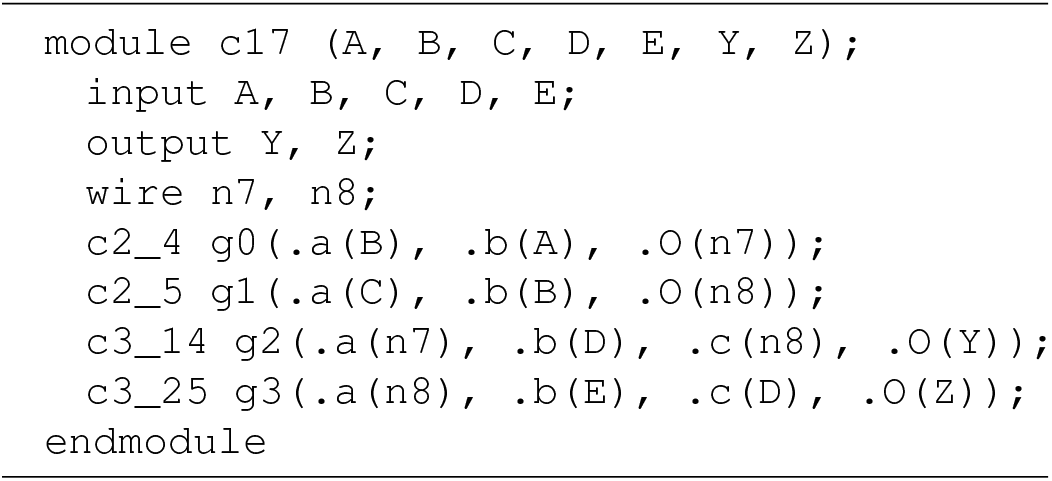
Synthesized c17 circuit in Verilog description.

Note that there can be more than one area-optimized circuit of a logic function. For comparison, in Fig. 11 we show another manually designed DNA implementation of c17 circuit whose area cost is 10 units as well. The corresponding DNA circuit is plotted in Fig. 12(B). Notice that the two circuits in Fig. 12 differ not only in their constituent logic gates, but also in their logic depths. The circuit of Fig. 12(A) is of two logic levels, whereas that of Fig. 12(B) is of three logic levels. There are six longest paths in the former circuit:

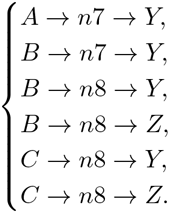

They involve a cascade of two logic gates. On the other hand, there are two longest paths in the latter circuit:

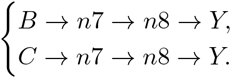

They involve a cascade of three logic gates. Although these two circuits have the same area cost, the circuit of Fig. 12(A) is preferred due to its better performance. In the experiments, we will synthesize circuits with area or performance optimized.

**Fig. 11.**
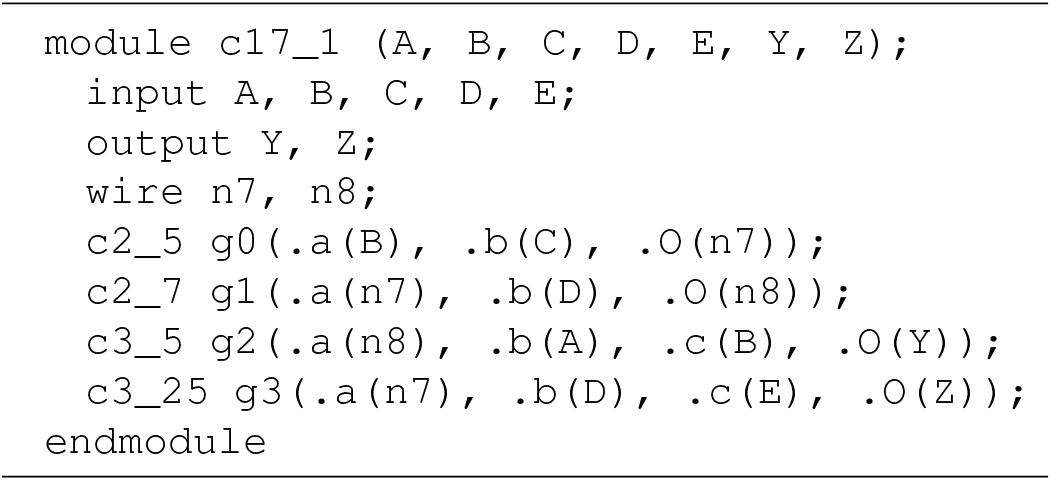
Manually designed c17 circuit in Verilog description.

**Fig. 12.**
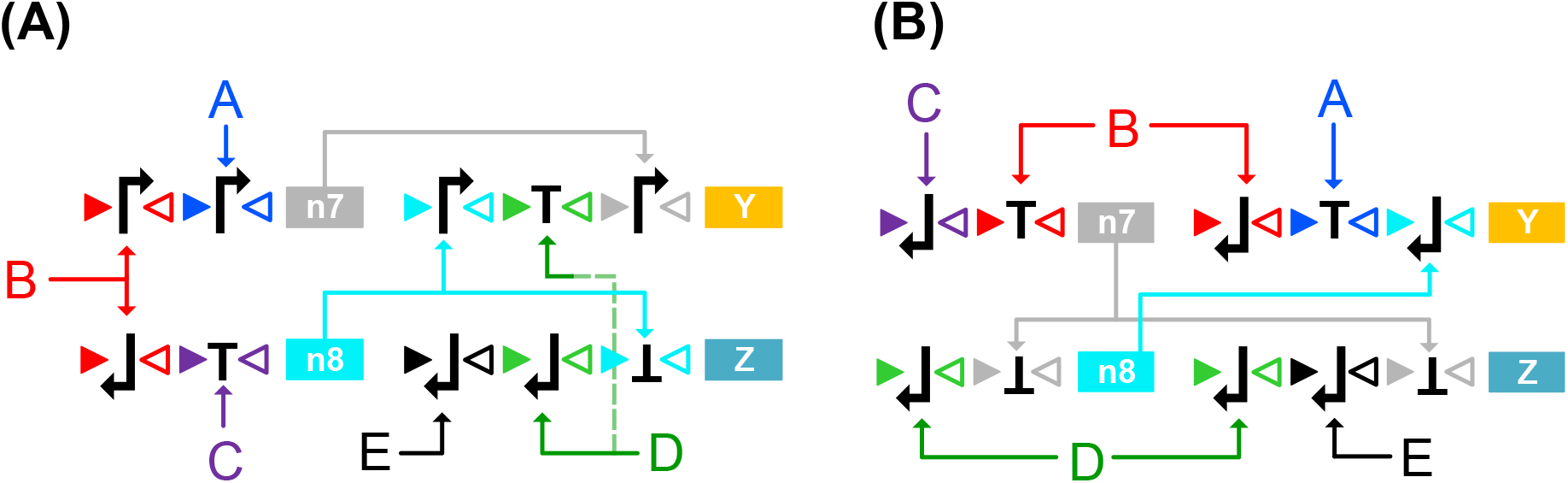
DNA circuit implementations of c17 benchmark circuit. (A) Implementation of the circuit in Fig. 10 synthesized by tool ABC; (B) Implementation of the circuit in Fig. 11 designed manually for comparison. In both (A) and (B), symbols *A*, *B*, *C*, *D*, and E indicate the recombinase inputs, the proteins encoded by the genes *Y* and *Z* are the outputs of the circuit, and the DNA gates encoding recombinases *n*_7_ and *n*_8_ and proteins *Y* and *Z* are the gates g0, g1, g2, and g3 in the modules c17 and c1_1, respectively.

## V. EXPERIMENTAL EVALUATION

To demonstrate the feasibility of the proposed synthesis flow, we experiment on other 67 ISCAS benchmark circuits using recombinase-based DNA gates. We expanded the library such that it includes all 684 DNA gates with decision list functions up to five inputs. In the library, the area cost of a gate is determined by the number of atomic terms, excluding the output gene, appearing in its corresponding 1g-wfs. We use a simple unit delay model for all the logic gates.

The experiment results of 54 (out of the 67) circuits are shown in Table II. The numbers of primary inputs/outputs, the number of inverters, and the number of logic gates (with the number of included buffers, if non-zero, reported in parentheses) are listed Columns 2, 3, and 4, respectively. The circuits were synthesized under two optimization settings: one for area optimization and the other for delay optimization. The results of area optimization are reported in Columns 5–7 and those of delay optimization are reported in Columns 8–10. For each synthesized circuit, its number of DNA gates, total area, and gate level are shown. In the naive implementations of benchmark circuits by simply converting the digital logic gates to the corresponding DNA gates, the total area of a DNA circuit can be roughly calculated as `“#inverter” + 2 × “#gate”`. Compared to the naive implementation, the circuits synthesized by ABC have much less area cost. Taking circuit b18 for example, we observe that the total area of the naive implementation is about **202110** which is much larger compared to the area **101870** of the area-optimized implementation and **105328** of the delay-optimized implementation. On the other hand, comparing area and delay optimized b18 circuits, delay optimization reduces the number of gate levels from **137** to **51** at cost of increasing area by **3500** units.

**TABLE II.** Results of Technology Mapping of ISCAS Benchmark Circuits

| circuit name | benchmark profile |  |  | area optimization |  |  | delay optimization |  |  |
| --- | --- | --- | --- | --- | --- | --- | --- | --- | --- |
|  | # PI/PO | # inverter | # gate (buffer) | # DNA gate | area | # level | # DNA gate | area | # level |
| b03 | 34/34 | 16 | 106 | 91 | 217 | 7 | 79 | 228 | 4 |
| b04 | 77/74 | 105 | 547 | 373 | 852 | 22 | 358 | 881 | 8 |
| b06 | 11/15 | 7 | 32 | 25 | 56 | 6 | 24 | 62 | 3 |
| b07 | 50/57 | 61 | 322 | 257 | 583 | 23 | 235 | 615 | 8 |
| b08 | 30/25 | 26 | 123 | 90 | 224 | 12 | 85 | 233 | 5 |
| b09 | 29/29 | 24 | 116 | 106 | 228 | 10 | 96 | 240 | 5 |
| b10 | 28/23 | 32 | 140 | 100 | 260 | 11 | 96 | 298 | 4 |
| b11 | 38/37 | 148 | 578 | 333 | 788 | 25 | 301 | 829 | 8 |
| b12 | 126/127 | 113 | 831 | 707 | 1648 | 15 | 673 | 1786 | 6 |
| b13 | 63/63 | 52 | 237 | 172 | 381 | 12 | 153 | 401 | 4 |
| b14 | 277/299 | 1531 | 8236 | 2851 | 6947 | 124 | 2791 | 7749 | 18 |
| b17 | 1452/1512 | 4474 | 26303 | 15344 | 37726 | 104 | 14802 | 39178 | 28 |
| b18 | 3357/3343 | 20372 | 90869 | 43018 | 101870 | 137 | 40277 | 105328 | 51 |
| b20 | 522/512 | 3068 | 16614 | 6119 | 14497 | 128 | 6111 | 16545 | 21 |
| b21 | 522/512 | 3089 | 16938 | 6173 | 14724 | 121 | 6147 | 16631 | 21 |
| b22 | 767/757 | 4491 | 24671 | 9302 | 22107 | 124 | 9286 | 24908 | 21 |
| c432 | 36/7 | 40 | 120 | 79 | 200 | 25 | 91 | 276 | 11 |
| c499 | 41/32 | 40 | 162 | 407 | 794 | 21 | 335 | 833 | 11 |
| c880 | 60/26 | 63 | 320 (26) | 234 | 530 | 26 | 208 | 553 | 8 |
| c1355 | 41/32 | 40 | 506 (32) | 394 | 781 | 19 | 328 | 878 | 10 |
| c1908 | 33/25 | 277 | 603 (162) | 336 | 690 | 28 | 271 | 736 | 13 |
| c2670 | 233/140 | 321 | 872 (196) | 409 | 956 | 19 | 400 | 1002 | 9 |
| c3540 | 50/22 | 490 | 1179 (223) | 566 | 1473 | 36 | 553 | 1649 | 14 |
| c5315 | 178/123 | 581 | 1726 (313) | 942 | 2202 | 25 | 908 | 2333 | 12 |
| c6288 | 32/32 | 32 | 2384 | 1825 | 3709 | 89 | 1502 | 3995 | 38 |
| c7552 | 207/108 | 876 | 2636 (534) | 1149 | 2496 | 59 | 1084 | 2754 | 11 |
| s208 | 19/10 | 35 | 61 | 39 | 100 | 8 | 39 | 105 | 3 |
| s298 | 17/20 | 44 | 75 | 55 | 125 | 7 | 52 | 138 | 3 |
| s344 | 24/26 | 59 | 101 | 82 | 178 | 11 | 67 | 175 | 4 |
| s349 | 24/26 | 57 | 104 | 84 | 179 | 11 | 67 | 175 | 4 |
| s382 | 24/27 | 59 | 99 | 78 | 172 | 8 | 70 | 191 | 3 |
| s386 | 13/13 | 41 | 118 | 71 | 186 | 7 | 61 | 195 | 3 |
| s400 | 24/27 | 56 | 106 | 80 | 173 | 9 | 76 | 220 | 3 |
| s420 | 35/18 | 74 | 122 | 79 | 202 | 11 | 72 | 196 | 4 |
| s444 | 24/27 | 62 | 119 | 75 | 169 | 9 | 74 | 210 | 3 |
| s510 | 25/13 | 32 | 179 | 116 | 311 | 8 | 102 | 324 | 4 |
| s526 | 24/27 | 52 | 141 | 88 | 202 | 11 | 79 | 223 | 3 |
| s641 | 54/43 | 272 | 107 | 94 | 217 | 17 | 82 | 232 | 6 |
| s713 | 54/42 | 254 | 139 | 90 | 212 | 16 | 85 | 237 | 6 |
| s820 | 23/24 | 33 | 256 | 130 | 353 | 8 | 129 | 394 | 4 |
| s832 | 23/24 | 25 | 262 | 132 | 358 | 9 | 135 | 406 | 4 |
| s838 | 67/34 | 149 | 241 | 163 | 415 | 12 | 142 | 398 | 5 |
| s1196 | 32/32 | 141 | 388 | 243 | 647 | 17 | 236 | 734 | 6 |
| s1238 | 32/32 | 80 | 428 | 278 | 734 | 17 | 259 | 790 | 7 |
| s1423 | 91/79 | 167 | 490 | 341 | 775 | 50 | 313 | 815 | 13 |
| s1488 | 14/25 | 103 | 550 | 299 | 820 | 12 | 272 | 910 | 4 |
| s1494 | 14/25 | 89 | 558 | 303 | 829 | 11 | 279 | 920 | 4 |
| s5378 | 214/228 | 1775 | 1004 | 844 | 1843 | 14 | 780 | 1849 | 7 |
| s9234 | 247/250 | 3570 | 2027 | 1065 | 2379 | 20 | 986 | 2442 | 9 |
| s13207 | 700/790 | 5378 | 2573 | 2006 | 4075 | 26 | 1818 | 4153 | 9 |
| s15850 | 611/684 | 6324 | 3448 | 2224 | 4946 | 35 | 2131 | 5018 | 16 |
| s35932 | 1763/2048 | 3861 | 12204 | 6776 | 14953 | 9 | 5565 | 14718 | 5 |
| s38417 | 1664/1742 | 13470 | 8709 | 6147 | 14319 | 23 | 5858 | 14551 | 10 |
| s38584 | 1464/1730 | 7805 | 11448 | 7066 | 16905 | 37 | 6243 | 16433 | 11 |

## VI. CONCLUSIONS

In this paper, we generalized the two-input recombinase-based DNA logic gates to multi-input cases. We formalized the syntax of recombinase-based logic gate construction, and obtained the Boolean function semantics of well-defined DNA sequences of recombinase-based logic gates. We also showed how to synthesize multi-level recombinase-based logic circuits using existing logic synthesis tools. Experimental results demonstrate the feasibility of our proposed methods. As recombinase-based logic circuits have been used in clinical biomarker detection, our results may automate complex recombinase-based circuit construction for advanced biomedical applications. With more and more evidence that DNA inversion can be mediated by genome editing tools such as the CRISPR/Cas9 system, we anticipate broad applications of recombinase-based logic gates in the future.

